# Metabolomic Profiling of Ashwagandha (*Withania somnifera*): Linking Bioactive Compounds to Functional Therapeutic Roles

**DOI:** 10.1101/2025.06.16.659832

**Authors:** Gaurav Tripathi, Vasundhra Bindal, Manisha Yadav, Neha Sharma, Sushmita Pandey, Babu Mathew, Nupur Sharma, Sanju Yadav, Vipul Sharma, Shvetank Sharma, Shiv Kumar Sarin, Jaswinder Singh Maras

**Author notes:** Corresponding Author: Correspondence: Dr Jaswinder Singh Maras, PhD Associate Professor, Department of Molecular and Cellular Medicine, Institute of Liver and Biliary Sciences, New Delhi-110070, Indià, <u> ext-24215</u> & Dr. Shiv K Sarin, MD, DM, D.Sc. (Hony.). Professor, Department of Hepatology, Institute of Liver and Biliary Sciences, New Delhi-110070, India. Equal contribution (Gaurav Tripathi, Vasundhra Bindal, Jaswinder Singh Maras). **Authors’ contribution:** JSM, SKS, SS and GT conceptualized the work. Drug profiling and experimental work and were helped by MY, NS, SY, BM, SP, VB, VS, and SP. Data analysis was performed by GT under the guidance of JSM. GT, SKS and JSM drafted the manuscript. This manuscript has been seen approved by all authors.

## Abstract

**Introduction:** *Withania somnifera* (Ashwagandha) is a widely used medicinal plant with established roles in traditional systems of medicine. Known for diverse pharmacological effects, including neuroprotection, hepatoprotection, enhancement of male fertility, and anti-inflammatory activity. Despite its longstanding use, a deeper understanding of its phytochemical composition and bioactive mechanisms is required to bridge traditional knowledge with modern therapeutic applications.

**Methods:** A comprehensive metabolomics profiling of *Withania somnifera* was performed to identify and quantify its bioactive secondary metabolites. Focus was placed on major classes such as alkaloids, phenolic compounds, and terpenoids. Known compounds were annotated and categorized based on their abundance, chemical class, and reported biological functions. Particular attention was given to compounds implicated in neuroprotection, liver health, reproductive support, and inflammation-modulation.

**Results:** The analysis revealed a rich but low-abundance of known metabolites. Alkaloids such as Gentianaine (8.44%) and Sauroxine (2.94%), and phenolics like Mulberrofuran A (14.58%) and Sulphuretin (9.61%), were among the most prominent. These compounds are associated with antioxidative, anti-inflammatory, and neuroregenerative activities. Terpenoids including Pinocarvone and Ergosterol contributed to hepatoprotective and anti-inflammatory properties. Compounds such as Gomphrenin-I, Sitosterol, Pfaffic acid, beta-Carotene and others (>3%) supported spermatogenesis, while Embelin and Kievitone enhanced oxidative stress response. Although most identified compounds were individually present in low percentages (<0.30%), their collective activity suggests synergistic effects.

**Conclusion:** Metabolomic analysis of Withania somnifera confirms high abundance of bioactive secondary metabolites that have hepatoprotective, neuroprotective, anti-oxidant and spermatogenesis-enhancing effects. These findings justify protective and beneficial effect of Withania in promoting stress relief, energy, and overall wellness.

**Graphical Abstract:** 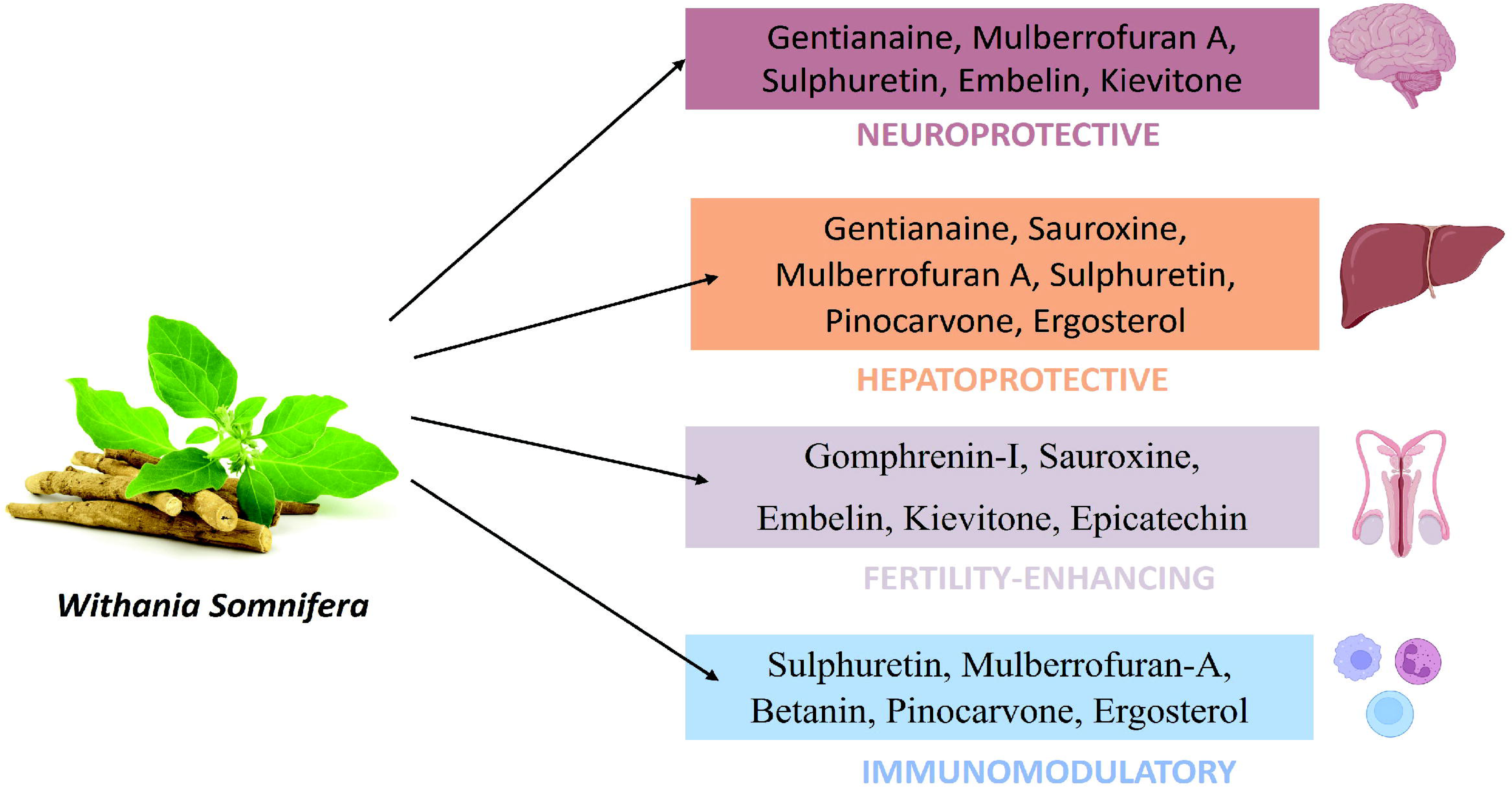

## Introduction

*Withania somnifera* commonly known as Ashwagandha, is a cornerstone of Ayurvedic medicine, traditionally employed to promote vitality, longevity, and resilience to stress. Modern pharmacological investigations have begun to substantiate these traditional claims, revealing a broad spectrum of bioactivities attributable to its diverse phytochemical constituents.^1^

Ashwagandha exhibits notable adaptogenic properties, aiding in stress mitigation by modulating the hypothalamic-pituitary-adrenal (HPA) axis and reducing cortisol levels.^2^ Its neuroprotective effects are evidenced by enhancements in cognitive function and memory, potentially through the upregulation of brain-derived neurotrophic factor (BDNF) and attenuation of oxidative stress.^3^ Furthermore, Ashwagandha demonstrates anxiolytic and antidepressant activities, which may be mediated via modulation of GABAergic and serotonergic pathways.^4^

In the realm of immunology, Ashwagandha has shown immunomodulatory effects, including the stimulation of lymphocyte proliferation, enhancement of natural killer (NK) cell activity, and increased phagocytic activity of macrophages. Its anti-inflammatory properties are attributed to the suppression of pro-inflammatory cytokines and inhibition of nuclear factor-kappa B (NF-κB) signalling pathways.^5^

Ashwagandha also contributes to metabolic health by improving insulin sensitivity and lipid profiles, thereby exhibiting potential in the management of metabolic disorders. Its hepatoprotective effects have been observed through the reduction of liver enzymes and protection against hepatotoxic agents.^6^

The herb’s influence extends to the gastrointestinal system, where it aids in maintaining gut integrity and modulating the microbiota, contributing to overall digestive health. Notably, Ashwagandha has been implicated in enhancing reproductive health, with studies reporting increased sperm count, motility, and testosterone levels in males, as well as improved sexual function in females.^7^

Despite these promising findings, the specific Secondary metabolites (bioactive compounds) responsible for known physiological effects are not well known, further their underlying mechanisms remain incompletely understood. Advances in metabolomics offer a comprehensive approach to elucidate the complex Secondary metabolites profile of Ashwagandha in relation with characterisation of its known physiological effects.

In this study, we employ untargeted high-resolution mass spectrometry-based metabolomics coupled to secondary metabolite screening using the plant metabolome data base (PMDB) to profile the secondary metabolite landscape of Ashwagandha. By integrating functional annotation to known physiological effects of Ashwagandha we wanted to identify secondary metabolites which provide hepatoprotective, neuroprotective, anti-oxidant and spermatogenesis-enhancing effects.

## Methods

To ensure robust and comprehensive metabolomic profiling, we utilized a meticulously optimized extraction and analytical pipeline. The *Withania somnifera* (Ashwagandha) root extract, sourced from AIIA, was initially solubilized using RIPA buffer and mechanically homogenized, followed by high-speed centrifugation. Both supernatant and pellet fractions underwent methanol-based protein precipitation and subsequent organic phase extraction. The samples were analyzed using ultra-high-performance liquid chromatography coupled with high-resolution Orbitrap mass spectrometry (UHPLC-HRMS), following reverse-phase separation through a C18 column. Internal and external standards were incorporated to validate analytical reproducibility across serial dilutions.^8–13^

Annotation using the PMDB phytochemical database revealed 1,020 compounds, spanning major classes of secondary metabolites—alkaloids, phenolics, and terpenoids—as well as six distinct plant hormone categories. This integrative and high-resolution profiling strategy not only highlights the chemotypic richness of Ashwagandha but also lays the groundwork for its application in evidence-based herbal medicine by linking its complex metabolome to diverse pharmacological activities.

The total compounds identified in *Withania somnifera* were initially annotated using the PMDB (Plant Metabolite Database). First, all annotated compounds were summed, and then the percentage abundance of each compound was calculated based on the relative intensity of their corresponding peaks. This approach allowed for an accurate determination of the phytochemical composition by expressing each compound’s contribution as a percentage of the total annotated metabolites detected in the sample(Figure 1A).

**Figure 1.**
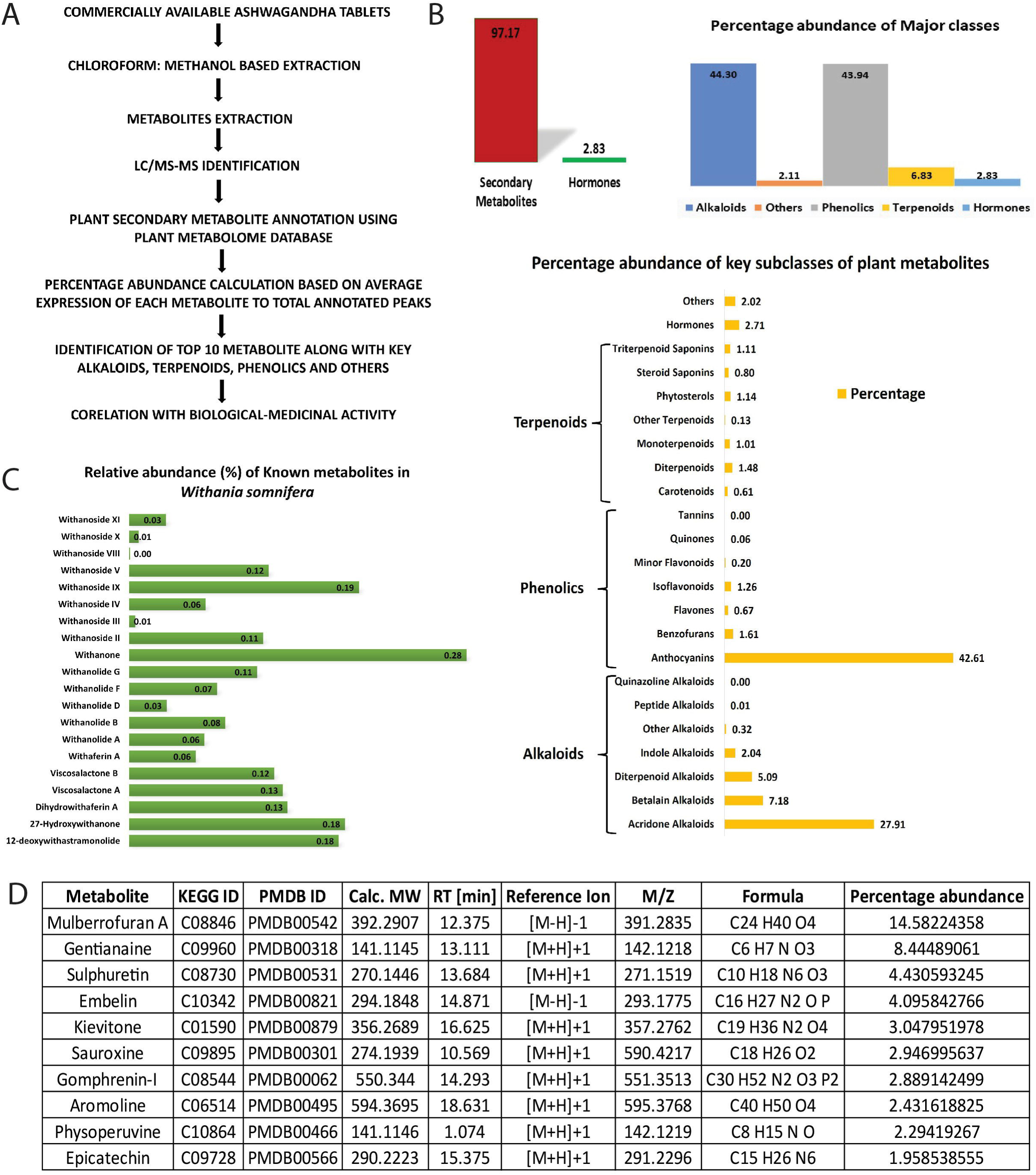
**A-** Study Design for the Proposed Research. **Figure 1B-** Percentage abundances of major and subclass categories of plant metabolites identified in *Withania somnifera.* **Figure 1C-** Percentage abundances of Known plant metabolites in *Withania somnifera.* **Figure 1D-** Table presenting the top 10 key metabolites identified in *Withania somnifera* using mass spectrometry, including m/z values, retention times, and related parameters.

## Results

### Distribution of Withanolides and Other Known Compounds in *Withania somnifera* Based on Percentage Abundance

The metabolic profiling of *Withania somnifera* reveals a diverse range of Withanolides and related compounds present in varying proportions. Among these, Withanone shows the highest percentage abundance at 0.28%, followed by Withanoside IX at 0.19%, and both 12[Deoxywithastramonolide and 27[Hydroxywithanone at 0.18% each. Other notable compounds include Dihydrowithaferin A and Viscosalactone A, each contributing 0.13%, while Viscosalactone B and Withanoside V are present at 0.12%. Withanolide G and Withanoside II show moderate abundance at 0.11%, whereas Withanolide F, Withanolide B, and Withanoside IV range between 0.06–0.08%. Compounds such as Withaferin A, Withanolide A, and Withanoside IV are detected at 0.06%, while Withanolide D, Withanoside XI, and Withanoside III are among the least abundant, each contributing below 0.03%. Notably, Withanoside VIII shows no detectable abundance in the profile. This distribution highlights the chemical complexity and variability of bioactive withanolides in *Withania somnifera* (Figure 1C).

### Major Phytochemical Classes in Ashwagandha Extract (Percentage Distribution)

The metabolomic profiling of *Withania somnifera* (Ashwagandha) revealed a diverse and functionally enriched phytochemical composition, dominated by phenolic and alkaloid subclasses. Anthocyanins emerged as the most abundant class, accounting for 42.61% of the total phytochemical content, followed by acridone alkaloids (27.91%), betaine alkaloids (7.18%), and diterpenoid alkaloids (5.09%). Other notable contributors included indole alkaloids (2.04%), plant-derived hormones (2.71%), Isoflavonoids (1.26%), diterpenoids (1.48%), triterpenoid saponins (1.11%), phytosterols (1.14%), and monoterpenoids (1.01%). Minor constituents such as quinazoline alkaloids, peptide alkaloids, quinones, and tannins were also detected, reflecting the chemo typic complexity of the extract. This broad-spectrum phytochemical distribution underscores Ashwagandha’s potential for multifaceted therapeutic activity, including neuroprotection, immunomodulation, metabolic regulation, and redox homeostasis, aligning with its traditional use in adaptogenic and rejuvenative medicine(Figure 1B).

The top compounds found in Ashwagandha, along with their percentage abundance, include Mulberrofuran A at 14.58%, Gentianaine at 8.44%, and Sulphuretin at 4.43%. Other notable compounds include Embelin (4.10%), Kievitone (3.05%), and Sauroxine (2.95%). Gomphrenin-I accounts for 2.89%, while Aromoline is present at 2.43%. Physoperuvine has a percentage abundance of 2.29%, and Epicatechin is found at 1.96%. Coronaridine and Afzelechin-(4α→8)-afzelechin are present at 1.95% and 1.73%, respectively. Additionally, Hildecarpin (1.70%), Poncirin (1.62%), and Muscapurpurin (1.42%) are observed. Other compounds include Rescinnamine (1.38%), Cadiamine (1.18%), Morindone (1.14%), Pseudobaptigenin (1.13%), and Gibberellin A15 at 1.06%(Figure 1D).

### Key alkaloids and subclasses in Ashwagandha

Out of the total secondary metabolites, which make up 97.17%, 44.30% are alkaloids. The alkaloid composition of the sample is primarily dominated by Acridone Alkaloids, contributing 27.91%. Betalain Alkaloids follow at 7.18%, while Diterpenoid Alkaloids are present at 5.09%. Indole Alkaloids account for 2.04%, and Other Alkaloids make up 0.32%. Peptide Alkaloids are present in trace amounts at 0.01%, and Quinazoline Alkaloids are the least abundant, representing just 0.002%(Figure 1B).

If the total alkaloids sum up to 100%, the key alkaloids contribute the following proportions: Gentianaine is the most abundant, representing 19.98% of the total alkaloids. Sauroxine and Gomphrenin-I follow, accounting for 6.97% and 6.83%, respectively. Aromoline and Physoperuvine are present at 5.75% and 5.43%, respectively, while Coronaridine contributes 4.60%. Other notable alkaloids include Muscapurpurin (3.36%), Rescinnamine (3.26%), and Cadiamine (2.79%). Betanin, Vasicinone, and Portulacaxanthin II make up 1.75%, 1.71%, and 1.71% of the total, respectively. Further, minor alkaloids such as Miraxanthin-II (1.45%), Akuammicine (1.26%), Vulgaxanthin-I (1.15%), and Ergosine (1.03%) contribute smaller proportions. Ergonovine, Akuammidine, Calebassone, and Garryine are present in the lowest amounts, each contributing under 1%(Figure 2A).

**Figure 2.**
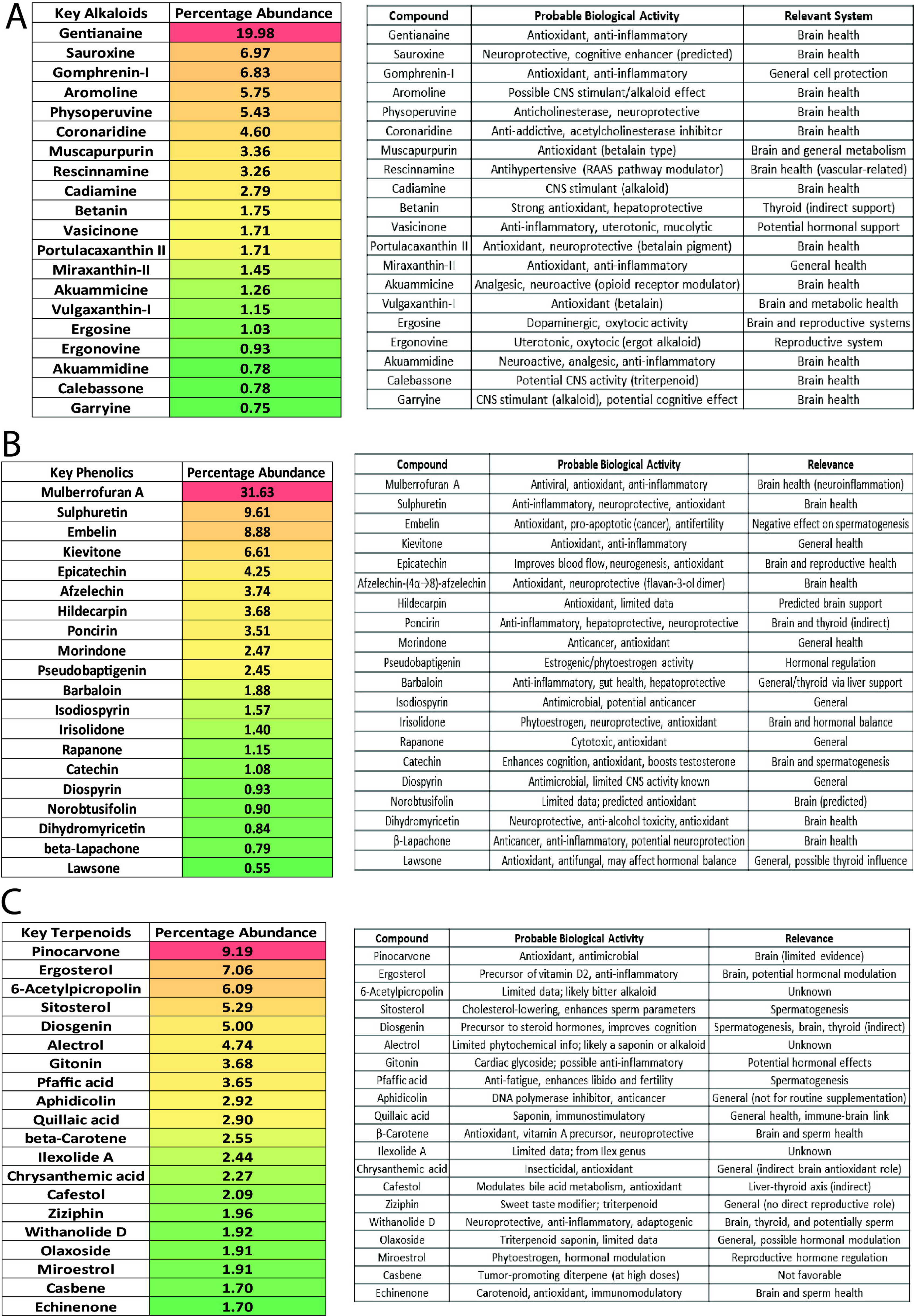
**A**- Percentage of key alkaloids detected in Withania somnifera using LC-MS/MS. **Figure 2B-** Percentage of key phenolics detected in Withania somnifera using LC-MS/MS. **Figure 2C-** Percentage of key terpenoids detected in Withania somnifera using LC-MS/MS.

**Figure 3.**
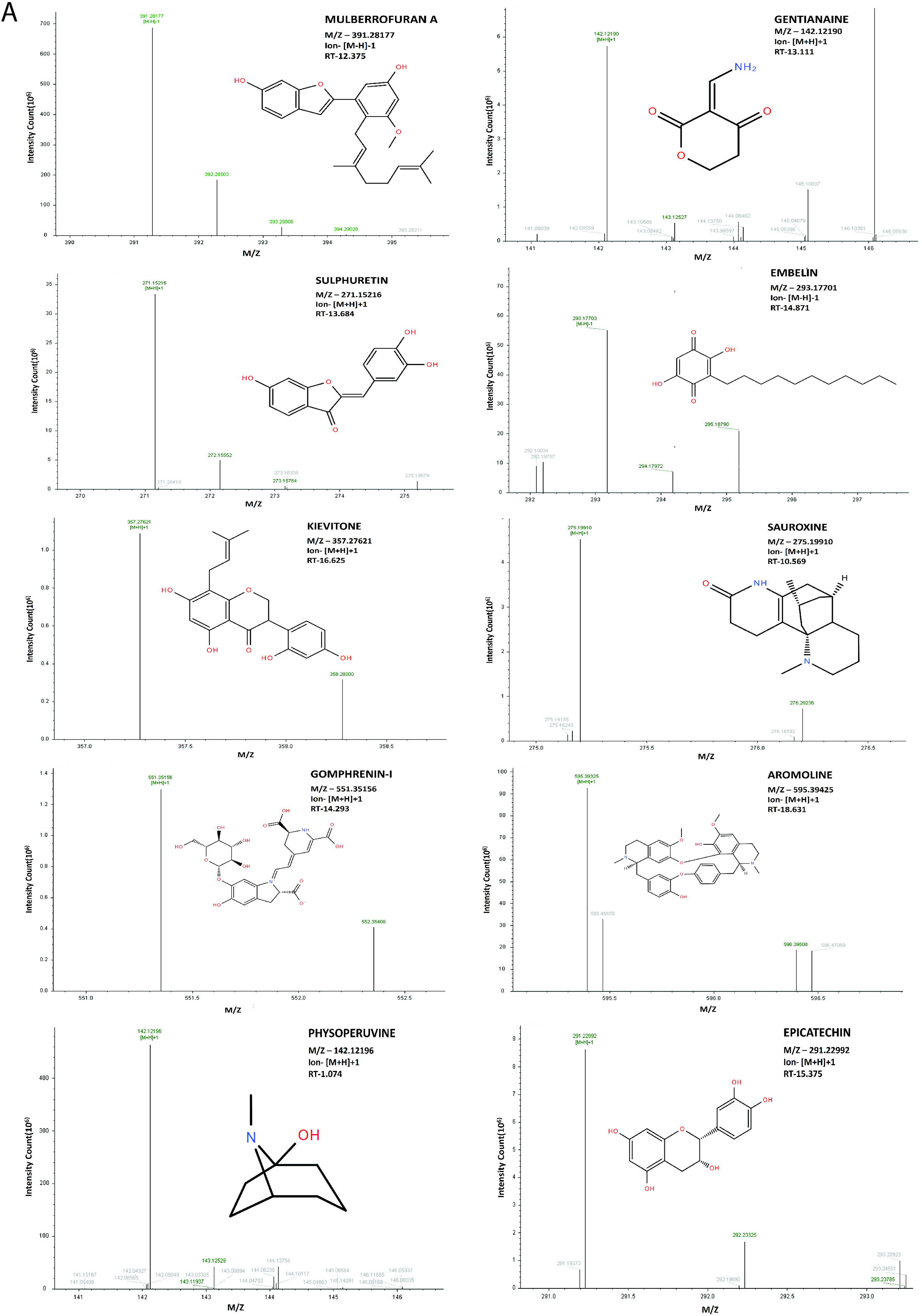
**A-** ion chromatogram of Top 10 key compounds identified in Withania sominifera.

### Key Phenolics and subclasses in Ashwagandha

Out of the total phenolics, anthocyanins make up the largest proportion at 42.61%. Other phenolic compounds include benzofurans (1.61%), Isoflavonoids (1.26%), and flavones (0.67%). Minor flavonoids contribute 0.20%, while quinones and tannins are present in very small amounts, contributing 0.06% and 0.004%, respectively. In total, the concentration of phenolics is 43.94%.

If the total phenolic concentration is 100%, the key phenolics contribute the following proportions:

Mulberrofuran A is the most abundant, representing 31.63% of the total phenolics. Sulphuretin and Embelin follow, making up 9.61% and 8.88%, respectively. Kievitone contributes 6.61%, while Epicatechin and Afzelechin-(4alpha->8)-afzelechin are present at 4.25% and 3.74%. Other notable phenolics include Hildecarpin (3.68%), Poncirin (3.51%), and Morindone (2.47%). Pseudobaptigenin contributes 2.45%, Barbaloin 1.88%, and Isodiospyrin 1.57%. Irisolidone, Rapanone, and Catechin make up 1.40%, 1.15%, and 1.08%, respectively. Minor phenolics such as Diospyrin (0.93%), Norobtusifolin (0.90%), Dihydromyricetin (0.84%), beta-Lapachone (0.79%), and Lawsone (0.55%) contribute smaller amounts to the overall concentration(Figure 2B).

### Key terpenoids and subclasses in Ashwagandha

Among the terpenoids, Diterpenoids have the highest abundance, accounting for 1.48%. This is followed by Phytosterols at 1.14% and Triterpenoid Saponins at 1.11%. Monoterpenoids contribute 1.01%, while Steroid Saponins make up 0.80%. Carotenoids represent 0.61%, and Other Terpenoids are present in the smallest proportion at 0.13%. The total concentration of terpenoids is 6.83%.(Figure 1B)

If the total terpenoids concentration is 100%, the key terpenoids contribute the following proportions:

Pinocarvone is the most abundant, making up 9.19% of the total terpenoids. Ergosterol follows with 7.06%, while 6-Acetylpicropolin contributes 6.09%. Sitosterol and Diosgenin account for 5.29% and 5.00%, respectively. Alectrol and Gitonin make up 4.74% and 3.68%, respectively. Pfaffic acid (3.65%) and Aphidicolin (2.92%) are also notable contributors. Other terpenoids include Quillaic acid (2.90%), beta-Carotene (2.55%), and Ilexolide A (2.44%). Chrysanthemic acid (2.27%), Cafestol (2.09%), and Ziziphin (1.96%) follow. Withanolide D, Olaxoside, and Miroestrol are present at 1.92%, 1.91%, and 1.91%, respectively. Casbene (1.70%) and Echinenone (1.70%) contribute the smallest amounts to the overall terpenoid concentration(Figure 2C).

### Physiological effect of ashwagandha metabolites

Withania somnifera (Ashwagandha) exerts wide-ranging physiological effects on the human body, largely due to its rich composition of bioactive compounds with diverse pharmacological properties. Key constituents such as Mulberrofuran A, Gentianaine, and Sulphuretin, which are present in relatively high abundance, contribute significantly to Ashwagandha’s anti-inflammatory, antioxidant, antiviral, and anticancer activities^14–16^. For instance, Mulberrofuran A (14.58%) demonstrates strong anti-inflammatory and antiviral effects, supporting immune modulation and protection against infections. Gentianaine (8.44%) offers neuroprotective and antibacterial benefits, helping safeguard the nervous system while combating microbial pathogens. Sulphuretin (4.43%) further enhances antioxidant defenses and exhibits anti-inflammatory and anticancer properties, promoting cellular health and reducing oxidative stress.

Other compounds such as Embelin and Kievitone also contribute to Ashwagandha’s antioxidant and antimicrobial effects, supporting overall immune function and potentially inhibiting tumor growth. Gomphrenin-I and Muscapurpurin provide hepatoprotective effects, protecting liver function by reducing inflammation and oxidative damage. The presence of Epicatechin and Afzelechin derivatives adds cardioprotective, antidiabetic, and anti-inflammatory benefits, which are crucial for metabolic and cardiovascular health^15, 17–19^.

Additionally, compounds like Sauroxine and Physoperuvine indicate potential central nervous system (CNS) stimulatory and anticholinergic activities, suggesting Ashwagandha’s role in modulating cognitive function and neurological health. Other constituents such as Poncirin and Pseudobaptigenin exhibit anti-inflammatory, antiallergic, and estrogenic activities, reflecting the adaptogen’s broad hormonal and immune modulatory effects^4, 20^.

Together, these bioactive molecules work synergistically to influence multiple physiological pathways—reducing inflammation, protecting vital organs, enhancing neurological function, and supporting cardiovascular and metabolic health—thereby underpinning Ashwagandha’s reputation as a powerful adaptogenic and therapeutic herb.

## Discussion

The diverse and functionally enriched phytochemical profile of *Withania somnifera* (Ashwagandha) underpins its wide-ranging therapeutic activities, including neuroprotection, hepatoprotection, promotion of spermatogenesis, and anti-inflammatory effects. The metabolomic profiling of Ashwagandha reveals a complex array of secondary metabolites, dominated by alkaloids and phenolic compounds, which together provide a solid basis for its traditional and emerging medical applications. In this study, we explored the percent distribution of major secondary metabolites and their association with multifaceted health benefits, including neuroprotection, hepatoprotection, spermatogenesis stimulation, and anti-inflammatory properties^1–6^.

The neuroprotective properties of Ashwagandha are primarily attributed to its rich alkaloid and phenolic content, particularly the presence of compounds such as Gentianaine, Mulberrofuran A, and Sulphuretin, which play a pivotal role in safeguarding the brain from oxidative stress and neuroinflammation. The alkaloid class, with Gentianaine as the most abundant compound (19.98% of alkaloids), has shown strong antioxidant capabilities, protecting neurons from free radical-induced damage^14–16^. Mulberrofuran A, the most abundant phenolic compound (31.63% of total phenolics), is also known for its neuroprotective effects, supporting neuronal growth and regeneration^14^. These compounds, along with Sulphuretin (9.61% of total phenolics), promote cellular health and may help prevent or delay the onset of neurodegenerative diseases such as Alzheimer’s and Parkinson’s. Furthermore, Embelin and Kievitone a polyphenol, which together account for over 10% of the phenolic content, exhibit neuroprotective properties by modulating oxidative stress pathways and promoting the survival of neuronal cells^17, 19^. The presence of Aromoline and Sauroxine within the alkaloid fraction, both of which have been implicated in neuroprotective mechanisms, adds further value to Ashwagandha’s potential in treating neurodegenerative conditions^20, 21^.

Ashwagandha also demonstrates considerable hepatoprotective properties, driven largely by its diverse alkaloid and phenolic content. The Acridone Alkaloids, contributing 27.91% of total alkaloids, are known to exert protective effects against liver toxicity by modulating oxidative stress and inflammation. Notably, Gentianaine and Sauroxine (6.97% and 6.83% of alkaloids, respectively) have been found to enhance liver function by reducing hepatocellular damage^15, 20^. Mulberrofuran A and Sulphuretin, key phenolic compounds, also play a significant role in liver protection, reducing lipid peroxidation and enhancing the antioxidant defense system, thereby mitigating liver damage from chronic diseases such as alcoholic liver disease and non-alcoholic fatty liver disease (NAFLD)^14, 16^. Additionally, Pinocarvone and Ergosterol, two important terpenoids, have been shown to exhibit hepatoprotective properties through their anti-inflammatory and antioxidative effects, helping to prevent liver fibrosis and improve detoxification pathways. These compounds, along with the overall rich composition of Ashwagandha, highlight its potential as a natural adjunct in the management of liver diseases^22, 23^.

The role of Ashwagandha in promoting male reproductive health is another area of considerable interest. Alkaloids such as Gomphrenin-I (6.83% of alkaloids) and Sauroxine are key players in stimulating testosterone production and supporting spermatogenesis. Gomphrenin-I has been shown to significantly enhance sperm count, motility, and morphology, making it a valuable compound in the treatment of male infertility^20, 24^. Additionally, Embelin, Kievitone, and Epicatechin—all of which are abundant in Ashwagandha—have antioxidant properties that protect sperm cells from oxidative damage, which is a major factor in infertility^17–19^. These compounds work synergistically to improve the health of the testes, increase sperm viability, and potentially boost overall male fertility.

Ashwagandha’s anti-inflammatory activity is one of its most significant therapeutic attributes. The phytochemicals present in Ashwagandha, particularly Sulphuretin, Mulberrofuran A, and Betanin, have demonstrated strong anti-inflammatory effects by modulating pro-inflammatory cytokines, such as TNF-α, IL-1β, and IL-6. Sulphuretin (9.61% of total phenolics) is particularly noteworthy for its potent ability to suppress inflammatory pathways in both the brain and peripheral tissues, offering protection against conditions like rheumatoid arthritis and chronic inflammation^14, 25^.

Pinocarvone and Ergosterol, key terpenoid compounds, further enhance the anti-inflammatory profile of Ashwagandha by reducing inflammatory cytokine release and promoting immune system modulation. These compounds work in tandem to alleviate symptoms associated with chronic inflammatory diseases, reducing the risk of systemic inflammation-related conditions such as cardiovascular diseases, diabetes, and neurodegenerative disorders^22, 23^.

In conclusion, the impressive phytochemical diversity of *Withania somnifera* (Ashwagandha) offers significant promise as a multifaceted therapeutic agent. Its neuroprotective, hepatoprotective, spermatogenesis-enhancing, and anti-inflammatory effects are deeply rooted in the synergistic actions of its bioactive compounds, such as Gentianaine, Mulberrofuran A, and Sulphuretin. As our understanding of these compounds and their molecular mechanisms continues to grow, Ashwagandha’s potential as an integrative treatment for a range of chronic conditions—from neurodegenerative diseases to infertility and liver disorders—becomes more evident.

Our method of identification of secondary metabolites using plant metabolome database offers a new paradigm for analysis of herbal products and characterising key secondary metabolites which may have important role in the pathophysiological effect exerted by the traditional Ayurveda formulation and drugs

